# New Cell Fate Potentials and Switching Kinetics Uncovered in a Classic Bistable Genetic Switch

**DOI:** 10.1101/215061

**Authors:** Xiaona Fang, Qiong Liu, Christopher Bohrer, Zach Hensel, Wei Han, Jin Wang, Jie Xiao

## Abstract

Bistable switches are common gene regulatory motifs directing two mutually exclusive gene expression states, and consequently distinct cell fates. Theoretical studies suggest that the simple circuitry of bistable switches is sufficient to encode more than two cell fates due to the non-equilibrium, heterogeneous cellular environment, allowing a high degree of adaptation and differentiation. However, new cell fates arising from a classic bistable switch without rewiring the circuitry have not been experimentally observed. By developing a new, dual single-molecule gene-expression reporting system in live *E. coli* cells, we investigated the expression dynamics of two mutually repressing transcription factors, CI and Cro, in the classic genetic switch of bacteriophage λ. We found that in addition to the two expected high-Cro and high-CI production states, there existed two new ones, in which neither CI nor Cro was produced, or both CI and Cro were produced. We constructed the corresponding potential landscape and mapped the transition kinetics between the four production states, providing insight into possible state-switching rates and paths. These findings uncover new cell fate potentials beyond the classical picture of λ switch, and open a new window to explore the genetic and environmental origins of the cell fate decision-making process in gene regulatory networks.

## Main Text

Cell fate decision-making is the process of a cell committing to a differentiated state in growth and development. The decision is often carried out by a select set of transcription factors (TFs), the expression and regulatory actions of which establish differentiated programs of gene expression (*1*). Bistable switches, which consist of two mutually repressing TFs, constitute a simple, widely conserved mechanism for cell fate determination. In a classic bistable switch, only one of the two mutually exclusive cell fates is possible, as only one of the two TFs can be stably expressed and maintained in a cell at any given time. Binary cell fate determination regulated by mutually repressing TFs has been observed in stem cell differentiation *(2, 3),* tissue development *(4, 5)*, bacterial competence development (6), bacterial conjugation (7), and phage infection *(8–12).* Multi-stability, the presence of more than two stable, mutually exclusive gene expression states, could arise from different circuitry configurations *(13–15).* In theory, multi-stability is possible even for classic bistable switches with weakened interactions between TF and genes in a stochastic, heterogeneous cellular environment *(16–19).* Here, we demonstrated experimentally the emergence of new expression states using the model bistable switch of the bacteriophage λ (20), offering new cell fate potentials beyond the classic genetic picture.

The λ switch is composed of two mutually repressive TFs, CI and Cro (Fig. 1A); the expression of CI but not Cro confers lysogenic growth, and the expression of Cro but not CI confers lytic growth. The λ switch has served as a paradigm for studying gene regulation and cell fate determination *(9–11, 20, 21),* but the real-time switching kinetics and paths between the two distinct, mutually exclusive gene expression states have not been elucidated experimentally. To achieve these goals, we developed a dual single-molecule gene expression reporting system to follow the stochastic expression dynamics of CI and Cro simultaneously in the same cells (Fig. 1B).

**Figure 1.**
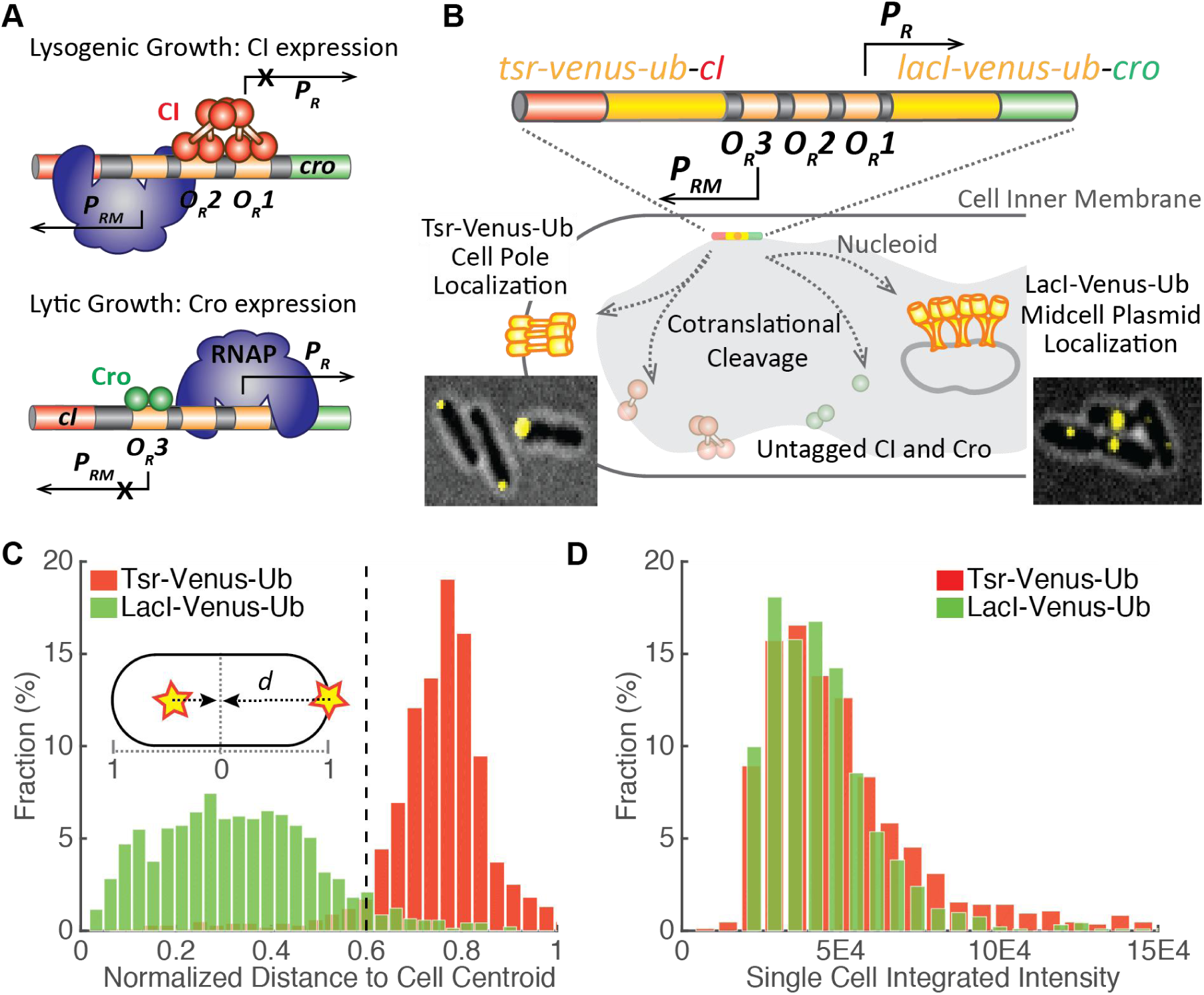
Validation of the dual single-molecule gene expression reporting system DuTrAc. (**A**) Schematic drawing of the two gene expression states of the λ genetic switch. In lysogenic growth, the λ repressor CI binds to its operator sites *O*_*R*_*1* and O_R_2 to stimulate its expression from promoter *P*_*RM*_, and at the same time shuts down Cro expression from promoter *P_R_.* In lytic growth, Cro binds to *O*_*R*_*3* to repress CI and turns on its own expression. (**B**) Schematic drawing of the DuTrAC system. The *λ* switch (colored bar) containing *tsr-venus-ub-cl* and *lacl-venus-ub-cro* is integrated into the chromosome at the *lac* operon locus. The expressed fusion polypeptide chain is cotranslationally cleaved by the constitutively expressed deubiquitinase UBP1 at the last residue of the Ub sequence, separating cell-pole-targeting Tsr-Venus-Ub from CI, and mid/quarter-cell-targeting LacI-Venus-Ub from Cro for single-molecule detection. CI and Cro are thus untagged and can bind respective operators to regulate gene expression. Representative cell images of pole- or midcell-localized Venus fluorescence spots were shown as insets. (**C**) Histograms of normalized distance (d) of Venus fluorescence spots to cell centroid (inset) in strain XF002 (red, expressing Tsr-Venus-Ub-CI only, fig. S1, Table S1) and XF003 (green, expressing LacI-Venus-Ub-CI only, fig. S1, Table S1) in the presence of the deubiquitinase UBP1. 92% of Tsr-Venus-Ub spots (n = 993 spots) localized at *d* ≥ 0.6 (dashed line); 95% of LacI-Venus-Ub spots (n = 1384 spots) localized at *d* < 0.6, suggesting that the threshold of *d* = 0.6 could be used to distinguish the identity of the fused protein. (**D**) Histograms of integrated fluorescence level of Tsr-Venus-Ub (red) and LacI-Venus-Ub (green) in individual XF002 and XF003 cells. The distributions and mean levels (4.8 ± 2.9 × 10^4^, n = 840 cells for XF002, and 4.2 ± 2.1 × 10^4^, n = 913 cells for XF003, μ ± s.d.) of Venus fluorescence in the two strains were indistinguishable from each other, indicating that both Tsr-Venus-Ub and LacI-Venus-Ub, despite of different mRNA and protein sequences, reported the expression levels of CI equivalently.

In the dual gene expression reporting system, we fused a fast-maturing yellow fluorescent protein variant, Venus (22) to one of two cellular localization tags, Tsr or LacI, to distinguish the production of CI and Cro in the same cell. The strategy of using two different subcellular localizations differs from previous studies using fluorescent proteins of different colors (23), and avoids the major disadvantage of temporal mismatches caused by different fluorescent protein maturation rates (e.g. ~1 hr for red fluorescent proteins such as mCherry *(24)* and ~5 min for Venus (22, *25–27)).* Tsr is a membrane protein that localizes rapidly and specifically to cell poles (28). LacI binds specifically to 256 *lacO* sites *(lacO^256^)* incorporated onto a multi-copy, mid/quarter cell-localizing RK2 plasmid pZZ6 (29). Because we can localize single fluorescent protein molecules with high precisions of 30—40 nm in live *E. coli* cells (27, *30),* we could distinguish between individual Tsr-Venus and LacI-Venus molecules based on their subcellular positions. Using a control strain XF004 expressing Tsr-Venus and LacI-mCherry independently (fig. S1, Table S1), we demonstrated that there was indeed minimal spatial overlap (~2%) between the two localization tags (fig. S2 and Movie S1).

Next, to distinguish the expression of CI and Cro in the same cell while avoiding possible disruptions of their functions due to the fluorescent protein fusion, we generated two translational fusion genes, *tsr-venus-ub-cI* and *lacI-venus-ub-cro,* and used the CoTrAC strategy (CoTranslational Activation by Cleavage, *(26, 27))* to cleave cotranslationally the Tsr-Venus-Ub or LacI-Venus-Ub reporter off from CI or Cro (Fig. 1B). This strategy ensures a 1:1 ratio in real-time between localized Venus reporter molecules and the fused CI or Cro molecules (or any other fused protein molecules, (27)). Using two control strains expressing only Tsr-Venus-Ub-CI (XF002) or LacI-Venus-Ub-CI (XF003), we verified that the cellular localization and fluorescence intensity of targeted Venus spots faithfully reported both the identity and expression level of CI (Fig. 1C and D, fig. S1, Movie S2 and S3). We named this new, dual gene expression reporting system DuTrAC (Dual coTranslational Activation by Cleavage).

To investigate the regulatory dynamics of CI and Cro in the λ switch using DuTrAC, we constructed strain XF204. We fused *tsr-venus-ub* to a temperature sensitive CI mutant (*cI*^857^ (31)), and *lacI-venus-ub* to *cro,* replacing the native *cI* and *cro* genes in the genetic switch, similar to what was previously described (fig. S1, Table S1) (27). We then integrated this circuit from *O*_*L*_ to the end of the *cro* gene into the chromosome of *E. coli* MG1655 strain at the *lac* operon locus (Fig. 1B). Hence, Tsr-Venus-Ub reports the expression of CI^857^, and LacI-Venus-Ub reports the expression of Cro. We used the temperature sensitive mutant CI^857^ (A66T, (31)) in place of wild-type (WT) CI^WT^ for the convenience of using temperature to tune the fraction of active CI. CI^857^ has normal DNA binding affinity and transcription regulation activity at the permissive temperature 30 ° C (32). At higher temperatures, an increasing fraction of CI^857^ becomes inactivated due to misfolding and subsequent degradation (*32*), and therefore Cro expression is switched on. We verified that the fusion of DuTrAC reporters to CI and Cro did not change the switching behavior of the genetic switch (fig. S3A). Furthermore, to examine the switching behavior across different protein expression levels, we generated two additional strains XF214 and XF224 (fig. S1, Table S1), in which the expression level of LacI-Venus-Ub-Cro was reduced in the order of [Cro]_XF224_ <[Cro]_XF214_ < [Cro]_XF204_. Using Western blotting we confirmed that the expression levels and switching behaviors of these strains were as expected (fig. S3B). Note that in the following experiments for simplicity we referred CI^857^ as CI.

To investigate the switching behavior of the modified λ switch, we first quantified the steady-state levels of CI and Cro in individual cells at different temperatures for strains XF204, XF214 and XF224 (Fig. 2, fig. S4, S5 and Table S2). We found that consistent with the typical bistable behavior, at a low temperature (30 ° C) cells had few Cro but predominately CI molecules (hereafter termed [L, H] for low-Cro and high-CI level, Fig. 2A, red curves); at a high temperature (37 ° C) the switch was flipped and cells predominately existed in high-Cro, low-CI level ([H, L], Fig. 2A, green curves). Interestingly, between the two extreme temperatures, we observed that a large population of cells had both CI and Cro [H, H] in the same cells at intermediate levels (Fig. 2A, black curves, fig. S5, Table S2). Reduced Cro levels in strain XF214 and XF224 did not abolish the presence of [H, H] population of cells, but shifted the temperature at which the percentage of this population of cells was the highest from 33 °C in XF204 to 34 °C in XF214 and to 36.5 °C in XF224 (Fig. 2, fig. S5, Table S2). A fourth population of cells having little CI or Cro ([L, L]) also existed at these temperatures. A few representative images of the four cell populations of strain XF224 at low, intermediate and high temperatures were shown in Fig. 2B. Note that under our experimental conditions (SOM), at each temperature cells were maintained at steady-state, i.e. on average the production of CI and Cro was balanced by their degradation, and hence their cellular concentrations, or steady-state levels, were not dependent on how many generations the cells have been grown in exponential cultures. Previous studies probing CI and Cro steady-state levels independently did not observe the presence of the two new populations of cells *(9,* 33).

**Figure 2.**
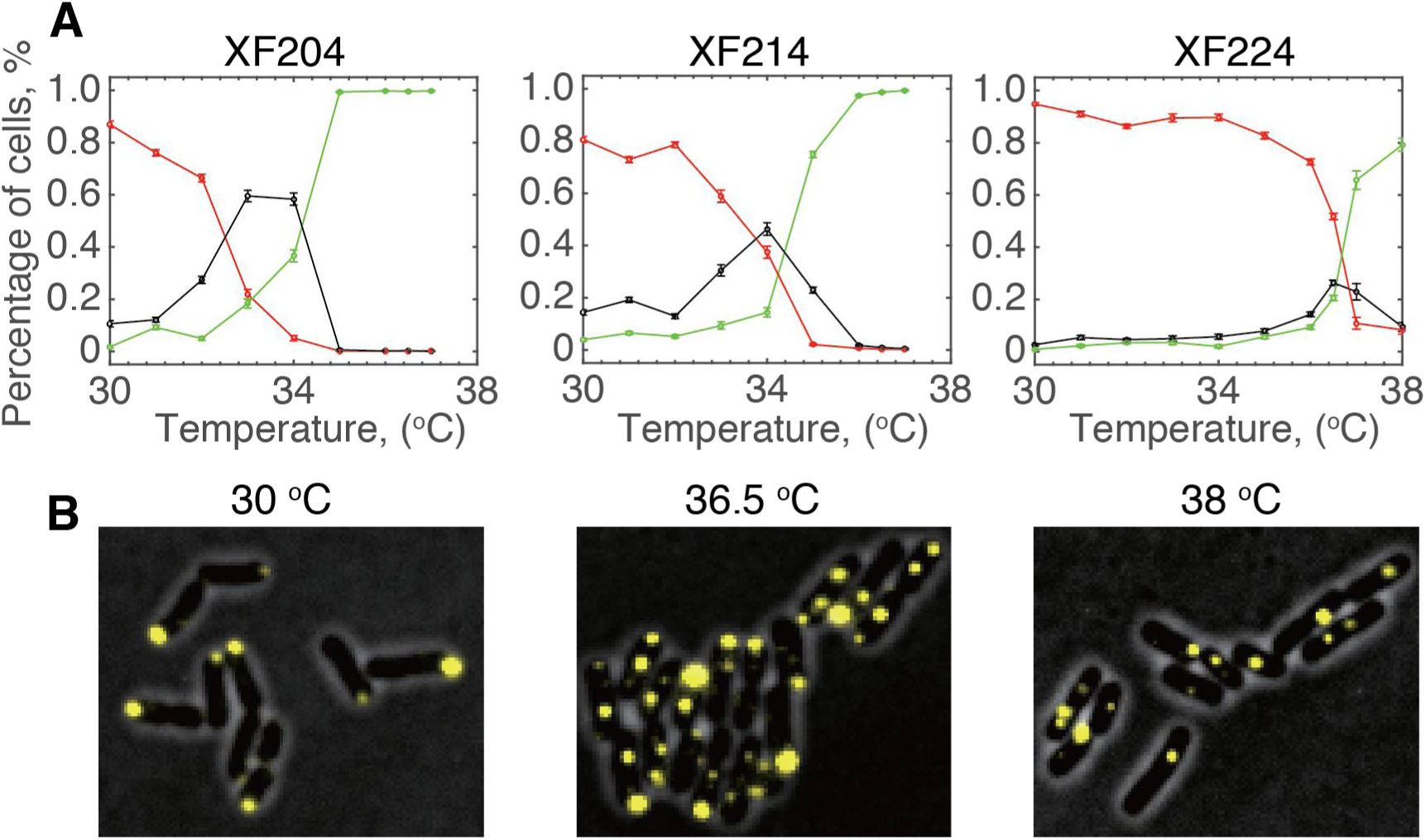
Steady-state levels of CI and Cro in strains XF204, XF214 and XF224 at different temperatures showed more than two expected cell populations. (**A**) Percentages of cells having CI only (red, copy number ratio *r =* CI/(CI+Cro) ≥ 0.8), Cro only (green, *r* ≤ 0.2), or both CI and Cro (black, 0.2 < *r* < 0.8) in strains XF204, XF214 and XF224 at different temperatures. See SOM for details. (B) Representative fluorescent images of XF224 cells showing CI expression (yellow pole-localized Tsr-Venus-Ub spots) and Cro expression (yellow quarter/midcell-localized LacI-Venus-Ub spots) at low, intermediate and high temperatures overlaid with phase-contrast cell images (gray).

The observation of cells having both CI and Cro at steady-state indicated that either these cells could switch between the two mutually exclusive steady-state levels of either CI or Cro within each other’s degradation time scales, and/or there existed a new expression state in which CI and Cro were expressed concurrently. The snapshot nature of the steady-state measurement could not distinguish these possibilities. In addition, the steady-state measurement of CI was complicated by the fact that at high temperatures an increasing population of CI^857^ becomes inactive; hence, the actual steady-state level of active CI (molecules/cell) is only proportional to the measured level of Tsr-Venus-Ub. These problems could be circumvented by following protein production in real time; the number of newly produced protein molecules per time unit directly reflects the promoter activity during that time without the convolution of any downstream processes. Therefore, we grew XF224 cells in a precision temperature-control chamber (T = 36.5 ± 0.1 °C over the length of the experiment of ~ 7 hours, fig. S6) on a microscope stage, and counted the number of newly produced CI and Cro molecules in individual cells every 5 minutes for multiple generations (SOM, fig. S7, Movie S4, S5, mean cell cycle time τ*=7* 1 ± 22 min, μ ± s.d., n = 457 cell cycles). We photobleached Venus molecules after each detection, so that new fluorescent molecules detected after five minutes of the dark interval were newly produced during the five minutes *(26, 27).* We chose strain XF224 because it had the lowest Cro steady-state levels compared to XF204 and XF214 (fig. S3, S5), facilitating the accurate identification and counting of single Venus molecules in small *E. coli* cells (fig. S8).

In Fig. 3 we show four representative time traces of different XF224 cell lineages. We observed stochastic, anti-correlated production of CI and Cro (Fig. 3A to D, fig. S9); Intriguingly, in many time traces, we observed that there were periods of time in which neither CI or Cro was produced, or both were produced. The presence of the four production populations was evident when we plotted the 2D histogram of the number of CI and Cro molecules produced in each 5-min imaging interval for all time traces (Fig. 3E). In addition to the two expected populations of high-CI ([L, H]), and high-Cro ([H, L]) production states, there were two additional populations. One resided at [0, 0] where no CI or Cro was produced, and another centered at ~ 4 molecules for both CI and Cro, similar to the [L, L] and [H, H] populations we observed in steady-state measurements (Fig. 2). The 1D histograms of CI and Cro alone showed two-state distributions (Fig. 3E).

**Figure 3.**
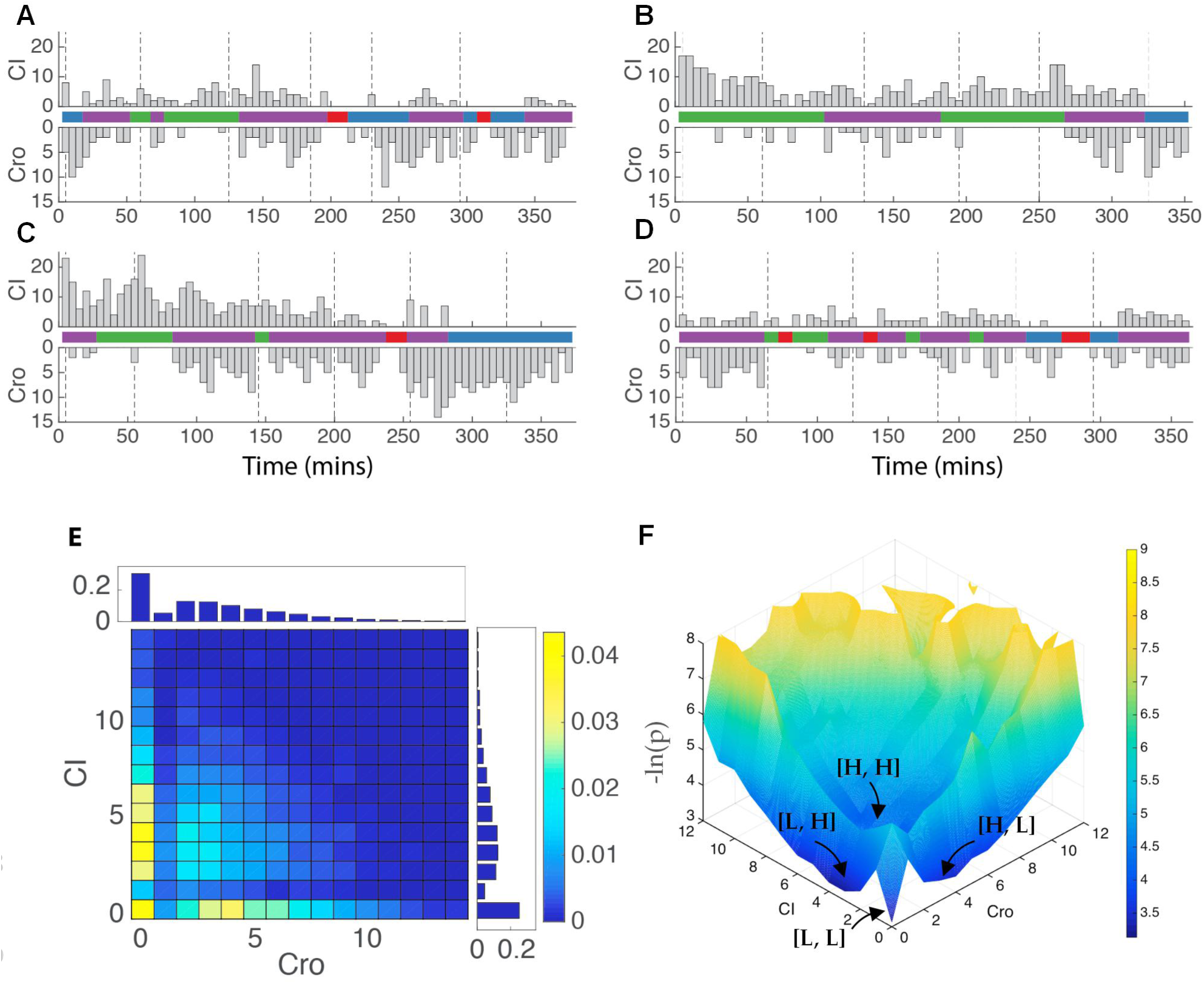
Real-time production time traces of CI and Cro in strain XF224 showed four distinct populations at 36.5 ° C. (**A** to **D**) Time traces of newly produced CI (top) and Cro (bottom) molecule numbers in the same cells in four representative XF224 cell lineages. The corresponding state-switching time trace identified by HMM was shown in the middle panel of each lineage, with blue corresponding to state [H, L] for high Cro and low CI production, green to [L, H], purple for [H, H] and red for [L, L]. The dashed vertical lines indicate cell division. (**E**) 2D histogram of produced CI and Cro protein molecules in individual cells measured at each 5-min frame in time-lapse experiments (n = 6453 frames from 94 time traces). Corresponding 1D histograms of CI and Cro are shown on the right and top of the 2D histograms respectively. Colors and scale bars indicate fractions of cells. (F) The potential landscape was calculated using the experimentally measured 2D histogram of CI and Cro expression numbers in every 5-min frame and interpolated.

We verified that the presence of the four populations was not caused by independent production of CI and Cro from two copies of the λ switch due to chromosome replication, because the four populations existed similarly in young cells where the chromosomal copy was one before replication (cell age <= 0.4, less than 40% of the cell cycle time, SOM, fig. S10A). Furthermore, single-molecule fluorescence *in-situ* hybridization (smFISH, Table S3) showed co-existence of *cI* and *cro* mRNA molecules in a significant population of cells (16.3 ± 0.7%, *cI* and *cro* mRNAs at 0.9 ± 0.03 and 0.6 ± 0.02 molecules per cell, μ ± s.e., n = 2627 cells), irrespective of cell ages (fig. S10B and Table S4). This result suggested that *cI* and *cro* mRNAs were produced within each other’s short lifetime window (~ 1.5 min, (34)). Co-existence of *cI* and *cro* mRNAs has also been previously observed in cells growing under a different growth condition (34). Finally, we verified that the stochastic maturation process of the Venus fluorophore only affected the spread, but not the presence, of each population in the 2D histogram (fig. S11). Taken together, these results suggested that in each 5-min time window, a cell could produce none, only one or the other, or both proteins.

Using the experimentally measured 2D distributions of CI and Cro production levels, we generated the corresponding potential landscape by calculating the negative logarithm of probabilities (Fig. 3F, SOM). There were clearly four basins, approximately around at [0, 0], [4, 0], [0, 4] and [4,4] for produced [Cro, CI] protein molecule numbers per 5 min (Fig. 3F). Interestingly, there was one central peak separating the four basins such that the barrier height between two opposite basins [L, L] and [H, H], or [H, L] and [L, H], was higher compared to that between two adjacent basins [L, L] and [L, H], or [L, L] and [H, L] (Fig. 3F). This type of landscape has not been previously observed for such a genetic circuitry, and suggested specific switching paths between the basins. For example, to switch from [H, L] to [L, H] the path going through the [L, L] or [H, H] basins would have higher probability than the path of switching directly between the two.

To quantitatively identify possible production states of CI and Cro corresponding to the observed basins in the potential landscape, and obtain the associated transition rates between these states, we used a modified Hidden Markov Model (HMM), which is commonly used in temporal pattern recognition *(35, 36).* We found that a four-state HMM ([L, L], [L, H], [H, L], and [H, H]) matched the observed 2D histogram of CI and Cro production the best (fig. S12). The mean production levels of Cro and CI of each state and the corresponding dwell time were summarized in Table 1 and fig. S13. Importantly, HMM allowed us to identify state-switching events in individual time traces (Fig. 3A to D, middle panels with colored bars) and hence the transition time constants (Fig. 4A, Table S5) (37). Similar results were observed using truncated time traces of only young cells (fig. S14, Table S5 and S6). Because the dynamics of a system is fully determined and described by its speed and the underlying kinetic processes (or paths), the transition time constants obtained here can be used to identify the most likely transition paths and the associated rates of switching between states, which has not been achieved before. For example, to switch from the [L, H] state to the [H, L] state, the most likely path is to go through the [H, H] state instead of directly switching. We can also determine the time it takes for switching by the times a cell spent on the two paths (from [L, H] to [H, H], and from [H, H] to [H, L]). This gives us insight into possible mechanisms underlying the kinetic processes in terms of the speed and the most likely paths, suggesting an unexpected kinetic route through [H,H] beyond direct switching between CI and Cro.

**Table 1:** Mean production levels of CI and Cro and the corresponding dwell time of each state identified by HMM in time-lapse experiments.

| State<br>[Cro, CI] | Cro<sup>a</sup><br>(molecules) | CI<sup>b</sup><br>(molecules) | n<br>(frames) | Dwell Time<sup>c</sup><br>(min) | n<sup>d</sup> (occurrence) |
| --- | --- | --- | --- | --- | --- |
| [L, L] | 0.0 ± 0.01 | 0.0 ± 0.0 | 139 | 17 ± 2 | 41 |
| [H, L] | 5.2 ± 0.10 | 0.0 ± 0.0 | 1173 | 36 ± 3 | 162 |
| [L, H] | 0.0 ± 0.0 | 4.7 ± 0.13 | 1451 | 27 ± 1 | 269 |
| [H, H] | 4.5 ± 0.04 | 4.7 ± 0.05 | 3069 | 47 ± 3 | 396 |

<sup>a</sup>, <sup>b</sup>, and <sup>c</sup>: Values were expressed as mean ± standard error.
<sup>d</sup>: Number of occurrence of each state in all time traces

**Figure 4.**
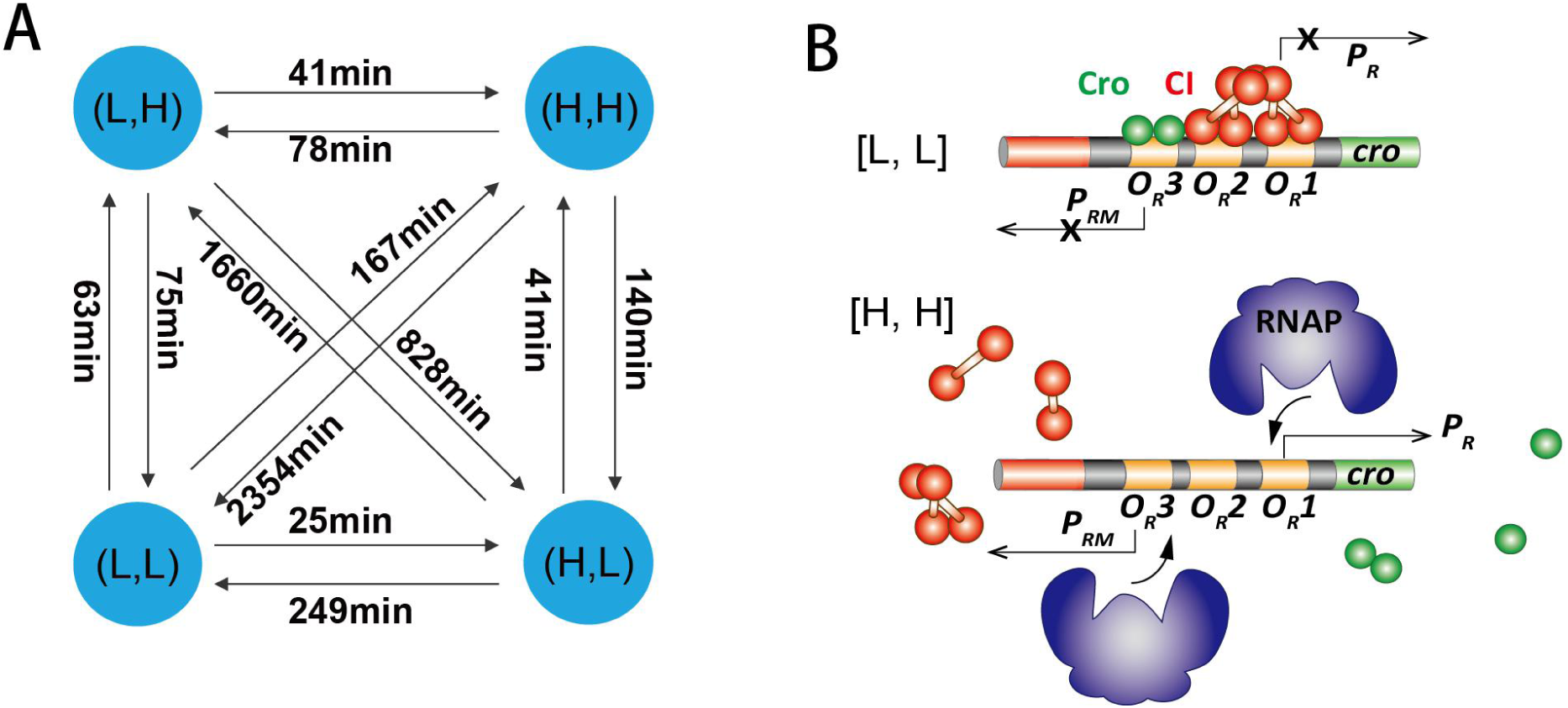
Transition time constants (A) and possible underlying molecular events (B) for the four stable production states observed in the λ switch. (**A**) Transition time constants were identified using a modified HMM analysis. Longer transition times indicate that direct switching between diagonal states is much less likely than that between side states. (**B**) The [L, L] states could arise due to the co-occupancy of the three operators by a combination of Cro and CI. The [H, H] state could arise from the slow association of Cro or CI to the operators, allowing RNAP to bind freely on *P*_*R*_ or *P*_*RM*_ to initiation transcription. Note that only these two simple possibilities were depicted here but other promoter configurations could also potentially lead to the [L, L] and [H, H] production states.

How does the λ switch generate two new production states [L, L] and [H, H], in addition to the two classic states [H, L] and [L, H]? Theoretical studies have shown that without changing the wiring configuration of a bistable switch, multistability can arise from weakened regulatory interactions, which impose fewer constraints on possible TF binding configurations *(16–19).* In eukaryotic cells, epigenetic phenomena such as histone modification and DNA methylation could reduce the binding rates of TFs to their targeting DNA sites, leading to longer time scale of gene regulation. In bacterial cells, low TF expression levels *(38)* and high levels of non-specific binding (39) can effectively slow down the binding of TFs to specific target sites, leading to weakened regulation. Such a regime in gene regulation is termed non-adiabatic, in contrast to the classic adiabatic regime, in which protein binding and unbinding are fast compared to the protein’s production and degradation time scales with rapid equilibration in a well-mixed environment *(17–19, 40).* The adiabatic regime of the λ switch under different experimental conditions has been elegantly demonstrated previously *(8, 41, 42).*

In our *λ* switch strain XF224, both CI and Cro are expressed at relatively low levels (fig. S15). Slow dissociation times (in the range of a minutes to hours *(43, 44))* and significant levels of nonspecific binding for both CI and Cro (> 70%, (39)) have been previously demonstrated. The [L, L] production state could emerge and remain stable when a combination of CI and Cro occupies all the operator sites, shutting down the production of CI and Cro simultaneously (Fig. 4B). When either CI or Cro dissociates, and the rebinding is slow, RNA polymerase (RNAP) can bind to the exposed *P*_*R*_ or *P*_*RM*_ promoters, resulting in the [H, L] or [L, H] production states. RNAP exists at a much higher level (~ 3000 molecules per cell under a similar growth condition, *(45))* compared to CI and Cro, and hence its binding rate would be significantly faster. When both CI and Cro dissociate from all three operators, which would occur at a much lower probability than only one of them dissociating, RNAP can initiate transcription from either one of the two promoters, resulting in the [H, H] production state (Fig. 4B). Consistent with this possibility, we previously measured the basal expression level of *P*_*RM*_ promoter in the absence of CI and Cro to be similar to the CI production level in the [H, H] state *(27).* In addition, *in vitro* studies have demonstrated that both *P*_*R*_ and *P*_*R*M_ promoters on the same λ switch can be occupied by two RNAP molecules simultaneously in the absence of CI and Cro *(46, 47).*

One important aspect of our time-lapse experiments, in contrast to an earlier experiment with a similar genetic network (8), is that we measured the production, but not cellular concentrations, of CI and Cro. Protein production rates directly measure promoter activities, which reflect promoter configurations with respect to TF and RNAP binding. In the adiabatic regime, new protein production instantaneously leads to changes in protein concentration and consequently protein binding rate; in the non-adiabatic regime, however, promoter activity does not necessarily follow immediate protein concentration changes. As such, for the same concentration state, different production states, and hence new cell fate potentials, could arise. The well known hysteresis effect of bistable switches (8) is likely a result of the non-adiabatic cellular environment in which protein binding/unbinding is slow— cells starting in one state have a tendency to stay in that state before switching to the other states even when the concentration of a critical protein has already changed.

Previous studies have shown that in the adiabatic regime, different wiring conditions of bistable switches could give rise to a maximum of three stable states *(13–15).* Here we showed that in the non-adiabatic regime, four stable protein production states can emerge from bistable switches without changing wiring configurations, with consequences in establishing new cell fates *(16–19, 48).* A living cell, in which non-equilibrium steady-state is the norm, could potentially utilize the non-adiabaticity to encode multiple cell fates even with a limited circuitry, hence allowing a high degree of adaptation and differentiation.

